# Beyond next-word prediction: hierarchical linguistic composition modulates LLM-brain alignment in time

**DOI:** 10.64898/2026.05.15.725490

**Authors:** Junyuan Zhao, Jonathan R. Brennan

## Abstract

The internal representations of large language models (LLMs) correlate, or “align”, with human neural activity during language comprehension. One view holds that this alignment reflects shared sensitivity to statistical patterns in LLMs and humans, while others hold that it reflects, at least in part, the emergence of shared linguistic representations in these systems. Here, we investigate whether hierarchical linguistic composition, a property believed to be fundamental to human language, modulates LLM-brain alignment. To this end, we manipulated syntax, compositional semantics, and associative semantics in English sentences that were presented to both an LLM and human participants during an electroencephalography (EEG) experiment. We matched linguistically manipulated stimuli in predictability, which allows us to tease apart alignment induced by linguistic structure from statistical factors. By comparing LLM-EEG alignment scores that were derived using a linear encoding model across predictability-matched conditions, we evaluate how linguistic manipulations modulate the alignment between human EEG reading data and contextual embeddings extracted word-by-word from the hidden layers of GPT2-XL. Three key patterns emerge: (1) increased alignment for word sequences with syntactic structure, (2) decreased alignment for sentences with compositional semantics, and (3) associative semantics does not modulate alignment. These observed linguistic modulations of LLM-EEG alignment take place above and beyond predictability. Our results indicate that associative semantics is encoded similarly by LLMs and the brain, as are at least some aspects of syntactic structure, while compositional semantics is more uniquely encoded in the human brain.

## 1. Introduction

The internal representations of large language models (LLMs) are shown to highly correlate with human brain activity during language comprehension. This so-called “LLM-brain alignment” has led to the proposal that LLMs may serve as models of human language processing in as much as they capture human-like comprehension through the mechanism of next-word prediction without prior knowledge specific to linguistic structure [1,2]. However, that two systems correlate does not necessarily entail that they deploy similar computations [3]. Critically, humans comprehend language through linguistic structures in a manner that dissociates from predictability: we can understand novel and unusual utterances which have not been encountered before so long as they are structurally well-formed [4]. Such processing involves the computation of hierarchical syntactic relations and compositional meaning [5–8]. It is therefore an open question what drives the alignment between LLMs and the brain. The alignment may reflect (1) shared sensitivity to statistical patterns and predictability alone [9], (2) convergent processing of linguistic structure that is learned from surface statistical patterns [10,11], or (3) coincidental convergence driven by the entanglement of linguistic structure and predictability in naturalistic data, which may not hold once predictability is controlled. In the current study, we manipulate three factors central to language understanding (syntax, compositional semantics, associative semantics) and investigate their contribution to the alignment between LLM hidden states and human electroencephalography (EEG) data. To separate alignment induced by linguistic structure from that induced by predictability, we match target items in terms of their LLM-derived next-word predictability. By dissociating structure from prediction, this study sharpens our understanding of the representations that may be shared, or unique, in human and artificial language processing systems.

### 1.1. LLM-brain alignment: findings, interpretations, and limitations

LLM-brain alignment quantifies the similarity between a computational language model and human neural signals. This typically involves finding a linear mapping between LLM hidden states and units of observation in human neuroimaging data (i.e., voxels in fMRI: see [1]; time-sensor cells in M/EEG: see [10]). Such a mapping from LLM hidden states to neural signals is an encoding model, and the prediction accuracy of such a model on held-out data yields a so-called “alignment score” [12]. In Schrimpf and colleagues’ foundational work [1], they found that the internal activation of LLMs captures nearly all explainable variance in human fMRI reading data. This correspondence between LLM representation and human fMRI increases with the specific model’s ability to predict the next word. Such alignment has been observed during listening, speaking, and reading across both hemodynamic and electrophysiological methods [10,13–15]. Furthermore, contextual representations extracted from the LLM captures cortical activity above and beyond static word representations even before a word is presented (e.g., [9]). The above findings are compatible with the fact that predictive processing is fundamental to both LLMs and the brain: The LLMs under discussion in the alignment literature are only trained to predict the next word in natural language data [16]; prediction is a key component for human language processing as well [17,18]. These observations motivate an account in which the capacity for successful next-word prediction alone may explain LLM-brain alignment.

However, prediction is only among many logical possibilities that lead to the observed alignment [19]. It is possible that any cognitive processes and task-relevant features that correlate with prediction may contribute to these observations. For example, Antonello and Huth [20] discuss how alignment is positively correlated with how well LLM internal representations generalize to other tasks, including part-of-speech tagging and translation. They suggest that, instead of predictability, LLMs and the brain align on generalizable task-specific features in language data that LLMs discover to solve downstream tasks. Even partial concordance on such tasks might lead to observed alignment patterns. This hypothesis intersects with arguments that human language processing diverges, in terms of task, with LLMs. In a computational simulation, Slaats and Martin [21] showed that LLMs do not necessarily separate predictability from grammaticality. The absence of separation here, where unusual instances of language become confounded with impossible instances of language, presents one challenge to the claim that LLMs are adequate cognitive models of human language processing [22,23] (but see [24]). Thus, our current understanding of LLM-brain alignment is limited in that on one hand, we do not know if features and processes other than prediction drive the observed alignment; on the other hand, it is unclear in the first place to what degree LLMs employ human-like processing.

As a window into the mechanistic understanding of LLM-alignment, we focus on the processing of hierarchical linguistic structure, a well-studied mechanism for sentence processing that is distinct from predictability. In the brain, hierarchical structure building is shown to capture human neural activity above and beyond predictability [6,25,26]. In LLMs, researchers have discovered latent syntactic structure by means of probing [27,28] and computational experiments [29–31] (cf. [32]). Recently, Kauf and colleagues [11] offered the first exploration into whether the alignment between LLMs and the brain may in fact be driven by syntactic structure. In that work, LLM-brain alignment to naturalistic language was modulated when the input to LLMs was perturbed. They found that instead of syntactic structure, the lexico-semantic content of language input most significantly modulates LLM-brain alignment. Yet, in this work they examined alignment only after manipulating language input to LLMs; they did not evaluate how alignment varied when the input to both LLMs and brains was manipulated. Thus, their findings offer only partial insights into the mechanisms driving alignment. Systematically studying how linguistic structure contributes to LLM-brain alignment requires simultaneously manipulating the structural properties of linguistic input in both LLMs and the brain. Furthermore, because statistical predictability captures variance along many linguistic dimensions [21,33–35], an adequate test for linguistic structure must also control for predictability.

### 1.2. The current study

In the current study, we aim to delineate the contributions of hierarchical linguistic structure to LLM-brain alignment while controlling for predictability. We study hierarchical linguistic structure via **syntax** and **compositional semantics**, along with **associative semantics**, a key non-structural component of linguistic meaning. Following previous work, predictability is quantified through next-word unexpectedness, or **surprisal**, of each word.

Syntax and compositional semantics are two related hierarchical features fundamental to language comprehension [8,36]. **Syntax** is the generative hierarchical system through which words combine into larger units of phrases and sentences that conform to symbolic and abstract rules [37–39]. Syntax is fundamental to complex meaning building, but on their own terms syntactically well-formed sentences do not necessitate a plausible interpretation just so long as its component words conform to structural constraints. For example, the sentence “*Colorless green ideas sleep furiously*” is as syntactically well-formed as “*Small colorful flowers bloom brightly*” despite the fact that the former has no sensible interpretation. Furthermore, researchers have found that syntactic computations may persist even in the absence of real words. The so-called “Jabberwocky” sentences, named after the poem by Lewis Carroll, contain pronounceable pseudo-words in lieu of meaningful lexical items yet still elicit the processing of sentence structure [40–42]. We thus test the effect of syntax on alignment by comparing sequences of pseudowords that do and do not form sentences.

**Compositional semantics** is the formal system through which word meanings are combined according to semantic rules under the constraints of syntactic structure [43,44]. Semantic composition does not yield a simple summation or intersection of word meanings but a complex output of a hierarchy of logico-semantic functions. For example, “*fake president*” cannot be interpreted by intersecting or adding together the meanings of *president* and *fake*; a “*fake president*” is not a president at all [8,43,44]. While even pseudoword sentences allow for some basic semantic operations, full-fledged meaning composition is only possible when real words combine in syntactically well-formed sentences. We thus operationalize compositional semantics as the joint contribution of real words and syntactic structure to alignment (detailed in Methods).

Finally, semantic composition must be dissociated from the **semantic associations** between words. These are non-hierarchical and reflect the conceptual co-activations that occur as part of language processing [45,46]. The role of associative semantics is established in the well-studied semantic priming phenomenon [47,48] as well as in cases where individuals infer sentence meaning despite impairment or disruptions in higher level aspects of sentence understanding [49,50].

We present English sentences that are manipulated along the above-mentioned linguistic dimensions to participants during EEG recording and also to the GPT2-XL language model [16]. Models of this class have been found to show robust alignment with neural signals in a wide variety of naturalistic and more controlled experiments [1,9,15]. Crucially, all target materials are matched in GPT2-XL surprisal to explicitly disentangle potential effects of linguistic structure from the effect of sequence predictability. LLM-brain alignment is quantified as the accuracy by which a ridge regression model predicts temporally unfolding EEG signal from word-by-word LLM embeddings. Alignment scores from different conditions are then compared to evaluate the effect of linguistic manipulations.

If next-word prediction exclusively drives LLM-brain alignment, namely that both the brain and LLM equally leverage statistical information, there will not be any difference in alignment value between conditions. This follows if next-word predictability is tracked in a similar way by both systems (Figure 1, left column). However, if alignment reflects one or more shared linguistic representations above-and-beyond predictability, we expect the following outcomes: If a factor is encoded both in LLM and the brain, word sequences with that factor will involve an additional facet of processing in both systems that goes above and beyond predictability, leading to an increase in LLM-brain alignment (Figure 1, middle column). This might be observed, for example, if LLMs represent at least some human-like latent syntactic structure [27,30]. If a factor is encoded uniquely in the brain, word sequences with that factor will engage additional processing only in the brain but not in LLMs. Such a discrepancy will yield reduced alignment (Figure 1, right column). Aside from alignment, we also test whether linguistic factors are encoded within internal LLM activations using probing methods [28] and whether these factors elicit decodable temporal-spatial patterns in EEG [51]. Because we compute time-resolved alignment values between EEG and LLM embeddings, we are able to pinpoint the difference induced by linguistic manipulations to specific time windows. This study advances our understanding of the shared and unique computations in LLM and human linguistic processing with precise temporal resolution.

**Figure 1.**
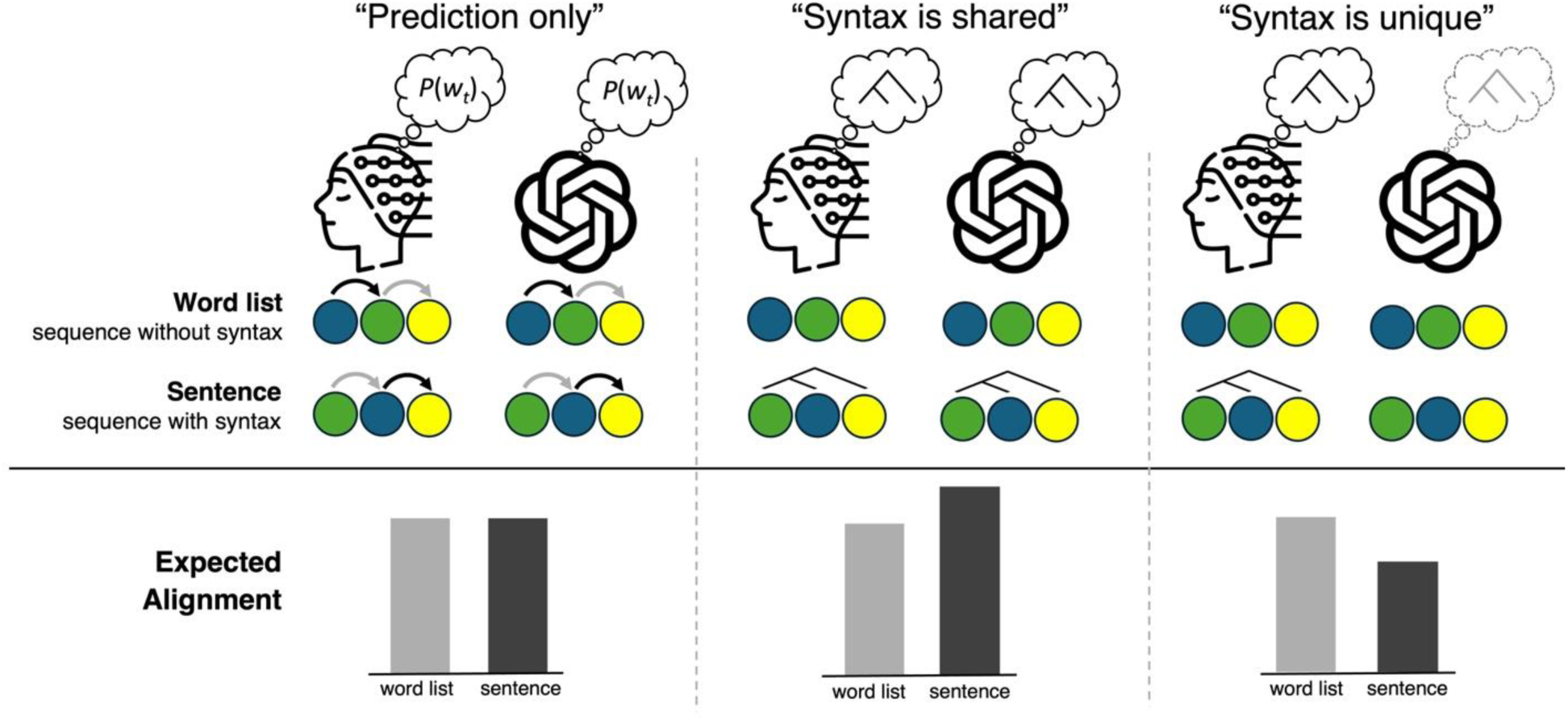
Predictions for LLM-EEG alignment. The two rows with colored circles represent hypothetical word sequences that differ in terms of a linguistic feature. Suppose the feature is syntax, the top row corresponds to word lists, and the bottom row corresponds to normal sentences. Note that here syntax is used as an example, while the reasoning outlined below may apply to any linguistic feature. Columns represent predictions for LLM and EEG alignment under different hypotheses. **The column on the left** represents the hypothesis that alignment is driven by shared encoding of next-word prediction. For both word lists and sentences, although the statistical pattern in the sequence may be different, the predictive mechanism used to track such pattern remains the same between LLM and the brain. Thus, we would expect no difference between LLM-brain alignment values calculated for word lists and sentences. **The column in the middle** represents the hypothesis that the brain and LLM both encode latent syntactic structures. Here, sentences differ from word lists in that they elicit structured representations. This additional shared process leads to increased alignment for sentences. **The column on the right** represents the hypothesis that only the brain encodes syntax. In this case, the brain and LLM similarly encode word lists. For sentences, only the brain processes syntax on top of single-word information, while LLM still processes the input as an unstructured word sequence. Such a divergence between LLM and the brain on sentences leads to a reduction in alignment under this hypothesis.

## 2. Methods

### 2.1. Participants

32 native speakers of English (10 men, 22 women, mean age = 21.0, SD = 1.4) participated in this study. All participants were right-handed with normal or corrected-to-normal vision. Participants were reimbursed for their time at 20 USD/hour. Written informed consent was obtained prior to the experiment. This study was approved by the Health Sciences and Behavioral Sciences Institutional Review Board at the University of Michigan (Protocol HUM00269683).

### 2.2. Design and materials

We presented participants and LLM with surprisal-matched English sentences that differed in linguistic details. We manipulated three linguistic factors: syntax, compositional semantics, and associative semantics. This was done by creating six conditions each comprised of 80 sentences in a parametric design: NORMAL, COLORLESS GREEN, JABBERWOCKY, SHUFFLED NORMAL, SHUFFLED COLORLESS, and SHUFFLED JABBERWOCKY. For details see Figure 2.

**Figure 2.**
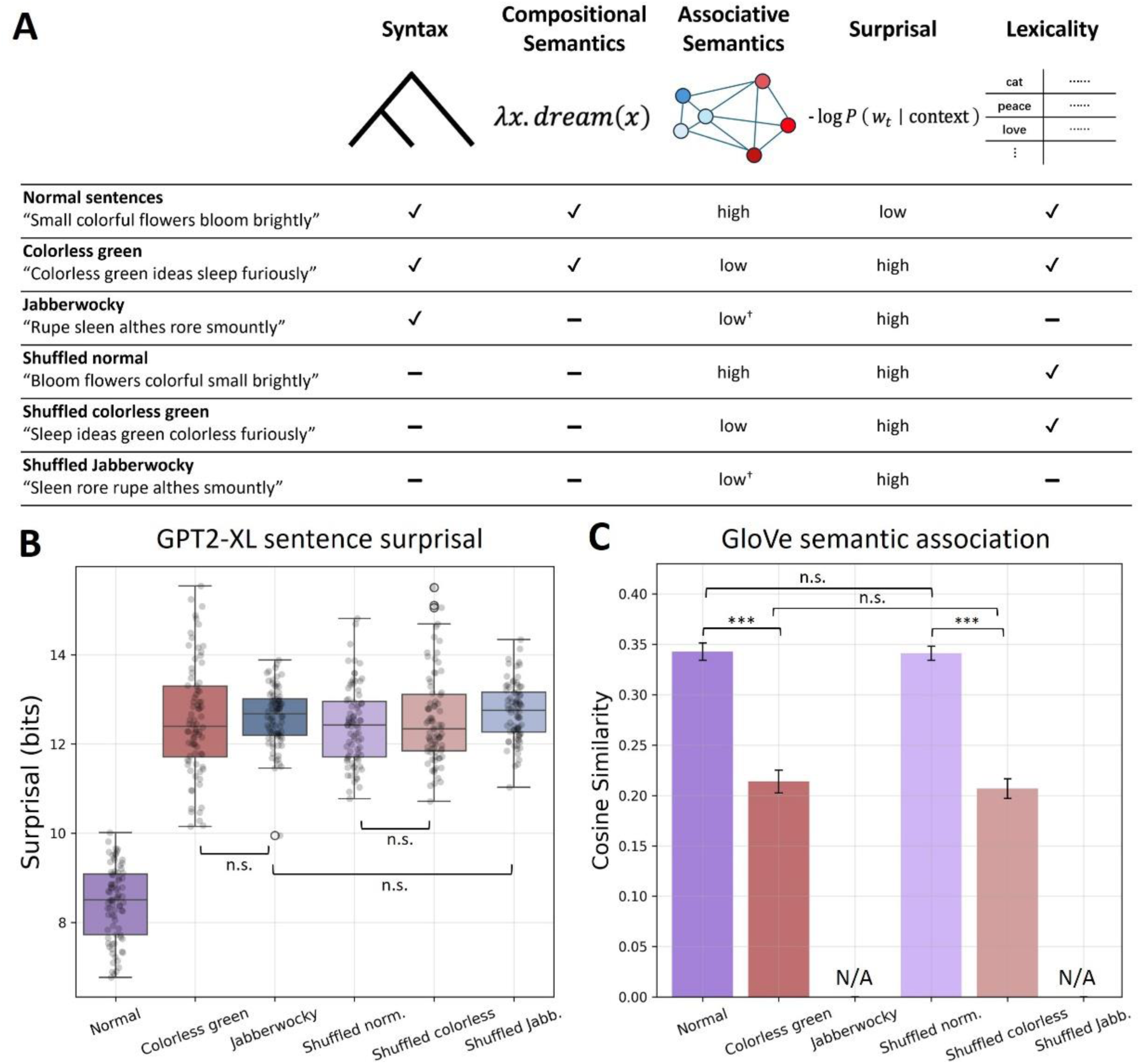
Experimental materials. (A) Design matrix with example sentences and factor annotations. B) Whole sentence GPT2-XL surprisal of all conditions. Sentence-level surprisal values were calculated by averaging over all tokens in a sentence. Contrasts of interest are tested with an independent samples *t*-test. (C) Mean pairwise cosine similarity of all conditions based on GloVe word embeddings (`glove.42b.300d`). This measure is not applicable to Jabberwocky and its shuffled counterpart because they do not have real lexical items. Effect of condition is tested with a two-way ANOVA on associative semantics and structure followed by Bonferroni-corrected pairwise post-hoc tests. Significance: * p < 0.05, ** p < 0.01, *** p < 0.001. †: Jabberwocky sentences have low semantic association because none of the pseudowords are lexically grounded, nor are they represented in GloVe embeddings.

**Figure 3.**
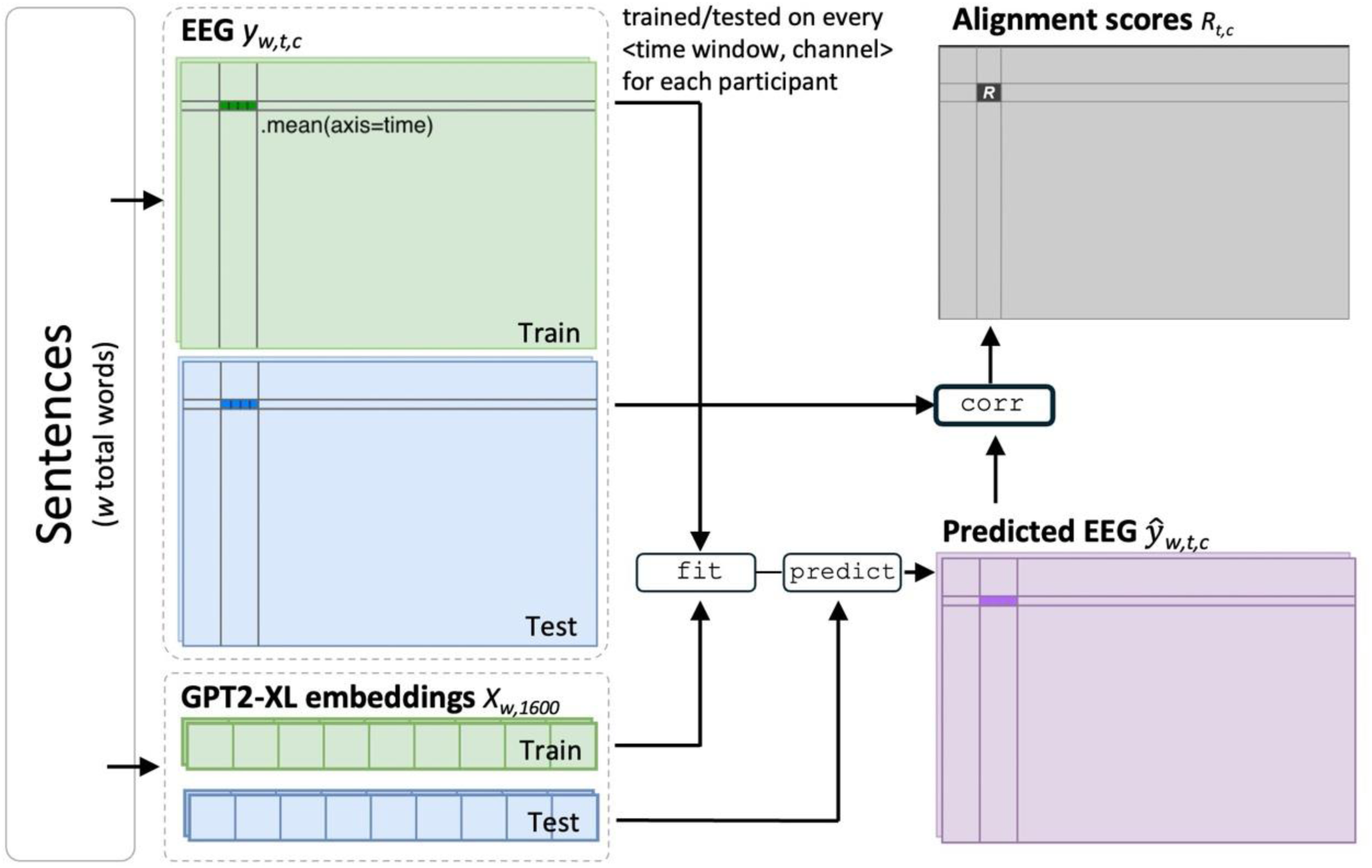
Illustration of the pipeline for computing LLM-brain alignment.

The NORMAL condition contains regular English sentences. The sentences used in this condition are five or eight words long, following two templates with a part-of-speech (PoS) order of either (1) *Adjective-Adjective-Noun-Verb-Adverb* (e.g., *Small colorful flowers bloom brightly*) or (2) *Determiner-Noun-Preposition-Determiner-Adjective-Noun-Adverb-Verb* (e.g., *These flowers in the small vase brightly bloom*). We generated NORMAL sentences with ChatGPT 4o (OpenAI, 2025; accessed on February 6, 2025) and manually identified those which were most natural. These sentences adhere to the syntactic and logical-semantic rules of English and have a coherent interpretation. Note that NORMAL is the only condition that is of low surprisal. This condition is used for stimuli generation, hyperparameter-tuning, and LLM layer selection.

COLORLESS GREEN sentences are syntactically well-formed. They license lexically grounded semantic composition [52,53], but lack a coherent interpretation [37,54]. COLORLESS GREEN sentences have weaker semantic association than in normal sentences. *F_assoc_*(1, 316) = 196.80, *p* < 0.001; *t*(158) = -9.00, *p* < 0.001; *F_struct_*(1, 316) = 0.20, *p* = 0.65. Items in this condition were created following the same PoS templates that were used for NORMAL sentences, except that we filled each word position with a word that is pseudo-randomly sampled from the SUBTLEX-US database [55], matching the PoS label in that position. Items that violate grammatical constraints (e.g., verb subcategorization, agreement) were manually excluded after sampling.

JABBERWOCKY sentences constitute pseudowords that follow the phonotactic rules of English but lack meaning. These pseudowords are combined with morphological cues to their grammatical properties and thus the structure of the sentence [42,56,57]. Consequently, JABBERWOCKY sentences have identical syntactic structure as NORMAL and COLORLESS GREEN sentences while lacking compositional semantics or associative semantics. We generated the pseudowords using the *Wuggy* Python package [58], which creates length-matched pseudowords by finding well-formed phonemic alternations.

Based on the above three conditions, we created three shuffled counterparts that do not have intact syntactic structure or compositional semantics. Despite the lack of syntax and compositional semantics, SHUFFLED NORMAL sentences retained higher semantic association than SHUFFLED COLORLESS sentences. *t*(158) = 11.08, *p* < 0.001. SHUFFLED JABBERWOCKY sentences contained no syntax, compositional semantics, or semantic association.

All conditions except NORMAL are matched in terms of last word surprisal. We first used the *minicons* Python package [59] to calculate last word surprisal of each sentence based on GPT2-XL. If a word is tokenized into multiple tokens, we take the average surprisal of sub-word tokens as the surprisal of the whole word. Then, we subsampled a larger set of candidate experiment sentences to yield conditions that are comparable in terms of last-token surprisal. The resulting subset does not show any significant difference in last word surprisal between the five high-surprisal conditions (*H*(4) = 5.01, *p* = 0.29, Kruskal-Wallis test). We further normed the stimuli in terms of whole sentence surprisal by averaging all tokens in a sentence. This analysis confirmed no significant difference between the critical contrasts (Figure 2B). The final experimental materials contained 6 conditions × 80 items/condition = 480 items. In addition, we included 320 grammatical filler sentences from the BLiMP dataset [60] to balance the total number of grammatical and ungrammatical sentences.

### 2.3. EEG procedure

Participants were comfortably seated approximately 100 centimeters from a computer screen in an isolated booth and were instructed to silently read sentences while minimizing movement and eye blinks. Sentences were presented word-by-word at the center of the screen in white text on a grey background using a rapid serial visual presentation (RSVP) protocol. Each sentence began with a fixation cross of 500 ms, followed by individual words for 300 ms and then a 300 ms blank-screen inter-stimulus interval. As an attention check, twenty percent of the trials were followed by a memory probe. Participants indicated whether they saw a specific word in the previous sentence by pressing one of two keys on the keyboard with their left or right index finger. The next trial began immediately after the key press or 500 ms after trials without a memory probe. Participants completed a short practice session to familiarize themselves with the task. The experiment session lasted approximately 1.5 hours. Participants were given the option of a short break approximately every ten minutes during EEG recording.

EEG signals were recorded using an ActiCAP 32-electrode system (Brain Products, Munich, Germany) distributed in a 10-20 layout. The EEG signal was amplified through a BrainAmp ActiCHamp+ DC amplifier, referenced online to a channel placed on the left mastoid, sampled at 500 Hz, and filtered with a passband of 0.1-200 Hz. The impedance of each electrode was kept below 25 kΩ by applying electrolyte gel prior to data recording.

### 2.4. EEG preprocessing

EEG signals were preprocessed and analyzed with the *MNE-python* package (version 1.10) [61] in a Python 3.11 environment. EEG data were re-referenced to the average of the left and right mastoids (TP9, TP10) and then filtered with a bandpass filter to 0.1-40 Hz. Then, the data were epoched from 0.2 s before word onset to 1.2 s post word onset for each word in a sentence. We used independent component analysis to remove artifacts resulting from eyeblinks and saccades (Jung et al., 2000). Trials with other artifacts such as muscle movement were automatically removed with the *autoreject* Python package (Jas et al., 2017; version 0.4.2). Participants with over 30% trials rejected were excluded from further analysis (epoch rejection rate before exclusion: median = 8.07%, range = 0 - 65.22%; after exclusion: median = 6.48%, range = 0 - 28.65%). Following this criterion, we excluded 4 participants and retained data from 28 participants.

### 2.5. Extracting representations from language models

We extracted contextual embedding representations of experiment items from GPT2-XL [16] hosted on *Hugging Face*. GPT2-XL is shown to have high correlation with human brain activity during naturalistic reading [1,62]. We fed the model one sentence at a time and extracted the 1600-dimension vector representation of each word from each of the 48 hidden layers. If a word is tokenized into multiple tokens, the average of the token-specific vectors is taken as the representation of the whole word. This procedure captures the model’s evolving representation as it incrementally processes sentences. Additionally, we used 300-dimension GloVe embeddings [63] of each word in a sentence as a non-contextualized control. GPT2-XL and GloVe embeddings were then reduced to 50 dimensions using PCA (following [9]) with over 90% of the variance retained. A baseline was created by replacing word-by-word embeddings with random 50-d vectors.

### 2.6. Quantifying LLM-brain alignment

LLM-brain alignment is quantified through the ability of a linear mapping model to predict neural response from LLM embeddings [1]. In this study, we obtained the alignment value by training and evaluating spatial-temporal ridge regression models.

First, for 80% of the sentences in a condition, we trained ridge regression models (implemented in the *himalaya* Python package [12], version 0.4.8) to find the linear transformation from GPT2-XL embeddings to EEG data. This is done at the word level, as EEG participants and GPT2-XL are presented with sentences word by word. Formally, the ridge regression can be expressed as an optimization problem for embeddings *X* and EEG data *y*:

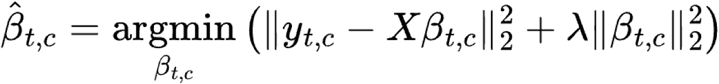

where a coefficient β is estimated for each time-channel pair (*t*, *c*) separately. To improve the signal-to-noise ratio, for each channel, we averaged EEG amplitude within 200-ms time windows that slide through time with a step size of 50 ms. The *l_2_* regularization parameter λ for each time-channel pair is optimized over a logarithmic range from 10^3^ to 10^10^ through ten-fold cross-validation. Then, we used the trained regression models to predict EEG response to words in the 20% held-out sentences. Again, this is done for each time-channel pair separately. Last, we computed cell-wise Pearson’s *r* between predicted EEG response *y*^ and the actual response *y* to unseen words. Namely, a separate correlation coefficient is computed for each time-channel pair over a total of *W* words from the sentences in a condition. The above procedure was cross-validated with ten contiguous data folds [64]. We interpret the average cross-validated correlation coefficient matrix *R* as the measure of LLM-brain alignment (“alignment score”). Alignment scores are computed within participants and for each condition separately. The embedding layer with the highest alignment score on normal sentences is used for further analysis.

### 2.7. Statistical analysis of LLM-brain alignment

Our parametric design enables planned comparisons that target the effects of specific linguistic factors. The effect of **syntax** is tested with a cluster-based permutation *t*-test between alignment scores from JABBERWOCKY and SHUFFLED JABBERWOCKY conditions [65] (implemented in the *eelbrain* Python package [66], version 0.39.0). This contrast allows us to target the effect of syntax without introducing lexical confounds [67,68]. The effect of **associative semantics** is tested through a permutation *t*-test between SHUFFLED NORMAL and SHUFFLED COLORLESS conditions, both of which contain no structure but differ in how words are conceptually related.

Because the factor of **compositional semantics** covaries with lexicality (see **Figure 2A**), we isolate this effect through an interaction in a two-way repeated measures cluster-based ANOVA that incorporates COLORLESS GREEN, JABBERWOCKY, SHUFFLED COLORLESS, and SHUFFLED JABBERWOCKY. In this ANOVA, compositional semantics is operationalized as lexically grounded syntactic composition, which means that any effect of compositional semantics will be captured by a difference-in-difference comparison expressed as (COLORLESS GREEN - JABBERWOCKY) – (SHUFFLED COLORLESS - SHUFFLED JABBERWOCKY). That is, a condition has (a potential effect of) compositional semantics only when there are both intact syntactic structure and real lexical items. To avoid confusion, we refer to the factor of syntax as (syntactic) structure in this statistical analysis. This is therefore an ANOVA for structure × lexicality.

### 2.8. LLM probing and EEG decoding

To further verify that our linguistic manipulations are meaningfully captured both in LLM and in the brain, we performed LLM probing and EEG decoding.

LLM probing is a classification-based method used to investigate if LLMs are sensitive to given linguistic features [28]. In this study, we trained logistic regression classifiers to distinguish the embeddings of sentence-final words drawn from a given contrast. Classification accuracy on a given contrast is computed for each GPT2-XL layer separately with 10-fold cross-validation and is interpreted as LLM’s sensitivity to that contrast.

We constructed three sets of contrasts of which JABBERWOCKY vs. SHUFFLED JABBERWOCKY targets LLM’s sensitivity to **syntax**, SHUFFLED NORMAL vs. SHUFFLED COLORLESS targets **associative semantics**, and comparing COLORLESS GREEN vs. JABBERWOCKY to its shuffled counterpart targets **compositional semantics** (following a similar logic detailed in section 2.7). The rest of the contrasts set up a broader context for interpreting LLM probing results, and these include NORMAL vs. SHUFFLED NORMAL, COLORLESS GREEN vs. SHUFFLED COLORLESS, NORMAL vs. JABBERWOCKY, NORMAL vs. COLORLESS GREEN, SHUFFLED NORMAL vs. SHUFFLED JABBERWOCKY, and SHUFFLED COLORLESS vs. SHUFFLED JABBERWOCKY. LLM probing accuracy is compared using FDR-corrected paired *t*-tests on all layers [69].

EEG decoding, similarly, is a classification method used to detect when the brain is sensitive to certain linguistic manipulations. We filtered EEG data between 1-50 Hz and created epochs from -1.0 to 3.6 s relative to sentence onset and trained a linear support vector machine EEG decoder within-participant on a given contrast, with a sliding window of 200 ms and a step size of 50 ms [70,71]. By decoding epochs that span 3.6 seconds (5 words) after sentence onset, we aim to capture how neural patterns evolve as sentences unfold. The decoder is evaluated with 5-fold cross-validation and yields an accuracy time series. The time series at the group level are then submitted to a temporal permutation *t*-test to discover time windows in which decoding accuracy of a given contrast was significantly above chance. For EEG decoding, we used the same contrasts as in LLM probing. EEG decoding accuracy is compared using a spatio-temporal cluster-based permutation *t*-test [65].

## 3. Results

### 3.1. Behavioral results

Participants achieved an overall accuracy of 90.2% (SD = 5.7%) on the memory probe task that followed 20% of the sentences. Data from two participants were excluded from this analysis due to an issue in behavioral data collection. Word recall accuracy is higher in low surprisal sentences (NORMAL) than high surprisal ones (the other conditions). *t*(25) = 3.85, *p* < 0.001 (Bonferroni corrected). We did not find any behavioral differences across the planned surprisal-matched contrasts laid out in section 2.7: syntax, *t*(25) = 0.71, *p* = 0.48; compositional semantics, *F*(1, 25) = 2.43, *p* = 0.13; associative semantics, *t*(25) = 0.66, *p* = 0.51.

### 3.2. LLM-brain alignment on normal sentences

This study asks whether LLM-brain alignment observed in language comprehension can be explained by predictability alone [1,9] and therefore we first identify the layer of highest alignment score based on normal sentences. This layer is then used for further comparisons that involve linguistic factors.

All hidden layers of GPT2-XL exhibited significant overall alignment with EEG data, with layer 17 being the most EEG-aligning layer that entered further analysis. The temporal dynamics of the alignment score is consistent across all layers: Hidden layers showed reliable pre-stimulus alignment with EEG data, followed by two peaks at around 200 ms and 850 ms (Figure 4). Pre-stimulus alignment is expected as our epochs include all words in each sentence, thus epochs extend into the presentation of the previous word. Layer-wise, alignment with EEG data increased in early layers (up to layer 17) and then shows a moderate drop after which it is stable until the final layers. Both the temporal evolution and the layer-wise pattern are consistent with alignment studies on data in other imaging modalities (MEG [10], ECoG [14]).

**Figure 4.**
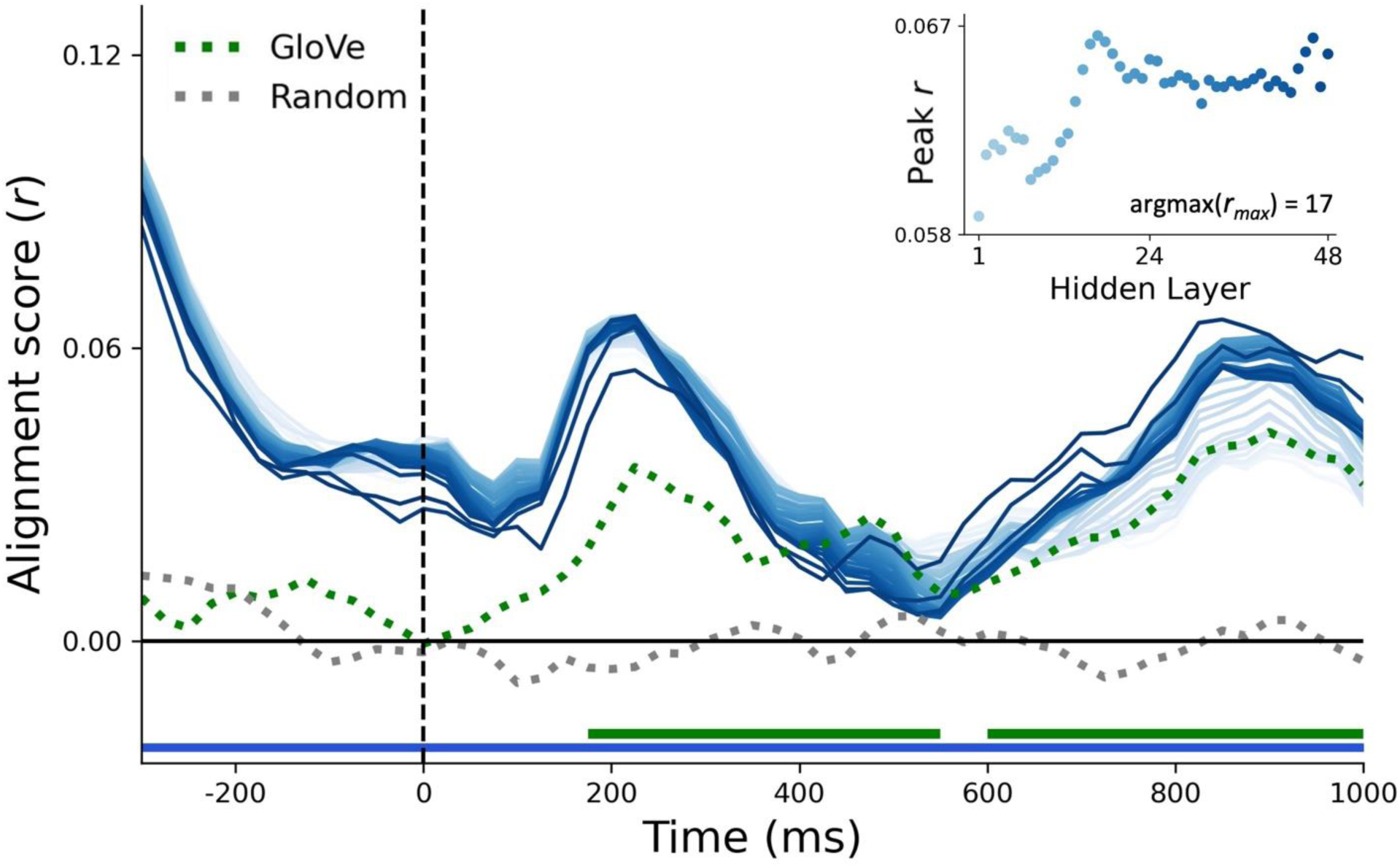
Alignment scores across embedding layers. Scores of GPT2-XL are colored with shades of blue, with darker shades indicating deeper layers. Layer-wise alignment is compared to random embeddings (dotted grey) and non-contextual GloVe word embeddings (dotted green). Figure inset shows peak post-stimulus alignment values of each layer. Colored bars above the x-axis indicate the significance of alignment scores relative to zero for GloVe (green) and for the average performance across GPT2-XL layers (blue).

GloVe word embeddings showed statistically reliable but lower alignment than most LLM hidden layers both at earlier (∼200 ms) and later (∼800 ms) time windows and they showed no significant pre-stimulus alignment. The absence of any pre-stimulus effect is expected for non-contextualized static word embeddings. Random embeddings, as a control, did not reach significance at any time point.

### 3.3. Linguistic modulation of LLM-brain alignment

Having established that layer 17 of the GPT2-XL large language model shows the best alignment to EEG signals and verifying that it outperforms static word embedding and random embedding controls, we compare the alignment score between conditions according to planned contrasts to test the effects of linguistic manipulations on LLM-brain alignment.

Alignment score increases in the presence of **syntax (Figure 5A)**. A cluster-based permutation t-test shows higher alignment score in JABBERWOCKY sentences compared to SHUFFLED JABBERWOCKY. *t_min_*(27) = 2.10*, p_cluster_* < 0.001. The significant cluster extends from 650 to 1,000 ms and is broadly distributed across the scalp.

**Figure 5.**
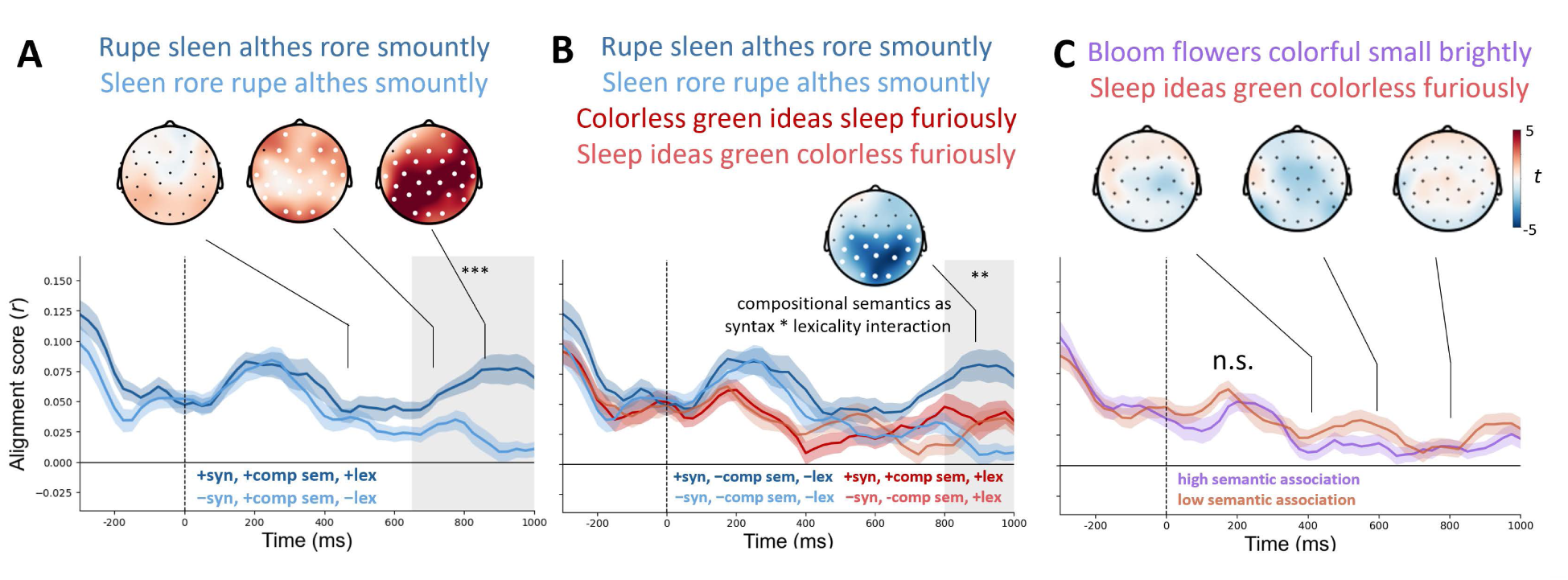
LLM-brain alignment of targeted comparisons. (A) Comparing JABBERWOCKY (dark blue) and SHUFFLED JABBERWOCKY (light blue) targets **syntax**. JABBERWOCKY shows higher alignment in later time-points with a broadly distributed scalp topography. (B) The interaction between JABBERWOCKY, SHUFFLED JABBERWOCKY, COLORLESS GREEN (dark red), and SHUFFLED COLORLESS (light red) targets **compositional semantics** and shows a reliable effect across posterior sensors after 800 ms. The effect is coded as a difference-in-difference: (COLORLESS GREEN - JABBERWOCKY) – (SHUFFLED COLORLESS - SHUFFLED JABBERWOCKY). (C) Comparing SHUFFLED COLORLESS and SHUFFLED NORMAL (light purple) targets **associative semantics**; no significant differences are observed. Shaded areas around alignment time series indicate 1 standard error of the mean. Time window of significant difference is marked in grey. Significance: * p < 0.05, ** p < 0.01, *** p < 0.001.

The presence of **compositional semantics** reduces LLM-brain alignment (**Figure 5B**). The cluster-based permutation ANOVA yielded significant main effects of structure (SHUFFLED vs. NON-SHUFFLED) and lexicality (JABBERWOCKY vs. COLORLESS GREEN) along with a reliable interaction effect. Alignment is higher in the presence of structure. This effect spanned 550-1000 ms and included all sensors. *F*_min_(1, 27) = 4.24, *p*_cluster_ < 0.001. Alignment is lower when a sentence contains real words. This effect is found in two clusters: The first cluster falls within 50-500 ms with a whole head topography. *F*_min_(1, 27) = 4.24, *p*_cluster_ < 0.001. The second cluster extends 825-1000 ms with a posterior scalp distribution. *F*_min_(1, 27) = 4.21, *p*_cluster_ = 0.047. Crucially, we found a significant interaction effect between structure and lexicality. *F*_min_(1, 27) = 4.23, *p*_cluster_ = 0.002. This interaction effect spans 800-1000 ms with a posterior scalp distribution. The consequent difference-in-difference test reveals that alignment is lower when a sentence contains both syntactic structure and real words, *t_min_*(27) = -5.12*, p_cluster_* = 0.002.

Lastly**, associative semantics** had no effect on LLM-brain alignment. *t*_max_(27) = 3.10, *p*_min_ = 0.10. Note that this null effect contrasts recent work by Kauf and colleagues [11] where the authors found a reduction in alignment when manipulating the lexico-semantic content of sentences. In contrast to the current study, Kauf et al. did not present semantically perturbed stimuli to human participants. As a result, any comparison between our results with theirs should be made with caution.

### 3.4. LLM probing results

LLM probing shows that all layers of GPT2-XL are sensitive to the linguistic contrasts we constructed, as evidenced by the above-chance decoding accuracy of the logistic classifier. Overall, we see a gradual increase in model’s sensitivity to linguistic variables across early layers and sensitivity plateaus in intermediate layers.

We further highlight the probing pattern at layer 17, which shows the highest brain-alignment score on normal sentences. This most brain-aligning layer shows reliable sensitivity to all conditions. Reliable sensitivity was observed at layer 17 on the minimal contrast between JABBERWOCKY and SHUFFLED JABBERWOCKY, which targets **syntax** (Figure 6A). As expected, the other non-minimal contrasts show greater accuracy.

**Figure 6.**
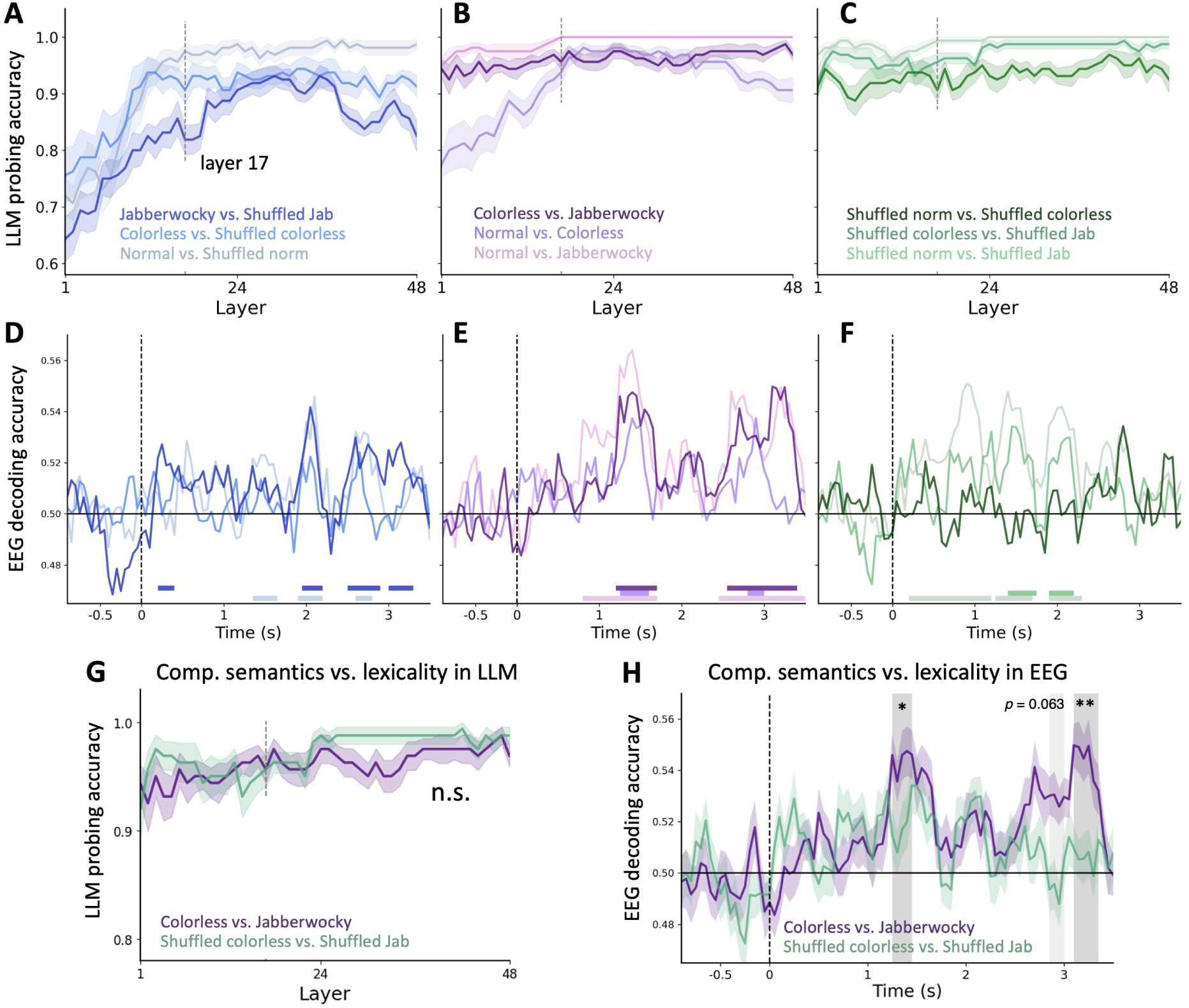
LLM probing and EEG decoding reveal sensitivity to linguistic manipulations in LLM and the brain. Top row: LLM probing; middle row: EEG decoding; lower row: lexicality confound analysis. (A,D) JABBERWOCKY vs. SHUFFLED JABBERWOCKY is the controlled contrast that explores both system’s sensitivity to syntax. (B,E) COLORLESS GREEN vs. JABBERWOCKY is the contrast that targets compositional semantics. (C,F) SHUFFLED NORMAL vs. SHUFFLED COLORLESS is the controlled contrast that targets associative semantics. The remaining contrasts are not minimal and are shown to provide a broader context for interpreting probing results. We additionally test whether lexicality alone explains the observed sensitivity to compositional semantics in LLM embeddings (panel G) and EEG signals (panel H). Dashed vertical lines in LLM probing panels mark layer 17, the layer with the highest alignment score on normal sentences which was used to compute LLM-brain alignment. Colored horizontal bars in EEG decoding panels indicate time windows of significantly above-chance decoding thresholded at α = 0.05. Grey shaded areas in panel H indicate time windows where compositional semantics is more reliably decoded in EEG signal. * p < 0.05, ** p < 0.01, *** p < 0.001.

This layer also shows sensitivity to contrasts related to **compositional semantics** (COLORLESS GREEN vs. JABBERWOCKY; SHUFFLED COLORLESS vs. SHUFFLED JABBERWOCKY). Crucially, there is no difference in LLM’s sensitivity to these pairwise comparisons across all layers |*t*(9)| < 2.50, *p_corrected_* > 0.61. The contrast COLORLESS GREEN vs. JABBERWOCKY entails additional difference in structure compared to SHUFFLED COLORLESS vs. SHUFFLED JABBERWOCKY, yet the LLM does not show differential sensitivity. This pattern indicates the LLM is more sensitive to whether these sentences contain real words (lexicality) than to whether the words additionally compose into larger units (structure). Taken together, an insensitivity to lexicality + structure suggests that LLM may not encode the kind of lexically rich linguistic composition that we term compositional semantics in the same way that humans do.

Regarding **associative semantics**, layer 17 shows sensitivity to SHUFFLED NORMAL vs. SHUFFLED COLORLESS. This sensitivity is equal to the comparison of SHUFFLED COLORLESS VS. SHUFFLED JABBERWOCKY and is lower than the non-minimal contrast of SHUFFLED NORMAL VS. SHUFFLED JABBERWOCKY (Figure 6C). Robust decoding accuracy in layer 17 for associate semantics ensures that the absence of a difference with respect to this factor in brain alignment (Figure 5C) cannot be explained as a lack of sensitivity to this comparison within the LLM alone.

### 3.5. EEG decoding results

We observe above-chance decoding for key linguistic contrasts as sentences unfold. This indicates at a coarse-grained level that our linguistic manipulations elicit stable differences in the signals under analysis. The principal contrast targeting **syntax** (JABBERWOCKY vs. SHUFFLED JABBERWOCKY) has the longest time window of above-chance performance among other pairs that also vary in structure (Figure 6D). The contrast of COLORLESS vs. JABBERWOCKY, which contributes to **compositional semantics**, has the highest decoding performance among pairwise contrasts between all conditions with syntactic structure (Figure 6E). The contrast targeting **associative semantics** (SHUFFLED NORMAL vs. SHUFFLED COLORLESS) is not decodable from this EEG signal, in contrast to other contrasts that co-vary in word association and lexicality (Figure 6F). The lack of decodability here may be an artifact of the sentence-level analysis which is less sensitive to semantic information distributed unevenly across these word lists. To test sensitivity to semantic associations at the word level, we fit temporal response functions (TRF [66,72]) between content word semantic dissimilarity [73] and epochs from these two conditions. This analysis shows that the semantic similarity between a word and the prior context explains additional variance in the EEG signal (detailed in Supporting Information S1).

We further compare decoding accuracy between COLORLESS GREEN vs. JABBERWOCKY and its shuffled counterpart (SHUFFLED COLORLESS vs. SHUFFLED JABBERWOCKY) to isolate the effect of **compositional semantics** from lexicality (Figure 6H). A cluster-based permutation test reveals that the difference in compositional semantics can be more reliably decoded in EEG than the difference in lexicality alone. The first cluster spans 1.25 – 1.45 s, *t_min_* = 2.20, *p_cluster_* = 0.047; a second, statistically marginal, cluster spans 2.85 – 3 s, *t_min_* = 2.63, *p_cluster_* = 0.063; the third cluster spans 3.1 – 3.35s, *t_min_* = 2.70, *p_cluster_* = 0.006. This finding complements the alignment difference induced by compositional semantics (Figure 5B) and indicates that the change in alignment score reflects the brain’s sensitivity to syntactically well-formed and lexically grounded sentences, a sensitivity that is absent in the LLM.

## 4. Discussion

In this study, we investigate how linguistic composition modulates the alignment between LLM internal representations and human EEG recorded while reading English sentences. Large language models are shown to align with the brain during language comprehension [1], yet it remains an open question whether this alignment reflects shared next-word prediction mechanisms [1,9], shared processing of latent hierarchical linguistic structure [10], or the coincidental entanglement of structure and predictability measures which are in fact encoded differently in LLMs and the brain [21]. To adjudicate between these possibilities, we dissociate the effect of linguistic composition from that of next-word predictability by controlling LLM-derived surprisal in linguistic manipulations. Manipulating syntax, compositional semantics, and associative semantics in both the GPT2-XL language model and human EEG recordings yielded three key findings: (1) The presence of basic syntactic structures increases the alignment between GPT2-XL and EEG. (2) The presence of compositional semantics reduces alignment. (3) LLM-EEG alignment is not modulated by associative semantics. The heterogeneous, rather than uniform, alignment patterns delineate the linguistic components on which an LLM and the brain converge and diverge. Because predictability is controlled across linguistic manipulations, these patterns highlight the modulation by hierarchical linguistic composition that is obscured by predictability measures or absent in prediction-only accounts.

Before turning to the effects of linguistic manipulations, we first validated our analytical pipeline on normal sentences. The alignment between LLM and EEG is quantified via a ridge regression encoding model that linearly maps the word-by-word embeddings of GPT2-XL to time-resolved EEG data recorded while participants read English sentences. All GPT2-XL hidden layers exhibit above-chance alignment with EEG signal on normal sentences and outperform non-contextualized GloVe embeddings. The alignment emerges before the onset of individual words and shows two peaks at approximately 200 and 800 ms post-onset, indicating that both systems leverage prior information to process current input [9,14,15]. Consistent with prior work [1,10], intermediate layers showed higher alignment than most early and late layers. Among intermediate layers, layer 17 exhibited the highest post-onset alignment at around 200 ms and was selected for further analysis. The temporal dynamics and layer-wise pattern of alignment on normal sentences validate our analytical pipeline and extend prior alignment findings to EEG.

Having established alignment that is consistent with the literature, we then ask how linguistic composition modulates alignment by comparing the magnitude of alignment values across our linguistic manipulations. Contrary to expectations from “prediction only” account (Figure 1, left column; [9]), the strength of alignment is not uniform across surprisal-matched linguistic manipulations. Rather, the three linguistic factors we manipulated show distinct modulations of alignment, pointing to the involvement of more fine-grained linguistic mechanisms in understanding the alignment, or misalignment thereof, between LLMs and the brain. We discuss each of these effects below.

### 4.1. Shared processing of basic syntactic structure in LLMs and the brain

We observed an increase in LLM-brain alignment for pseudoword sentences that contain **syntactic structure** (the JABBERWOCKY condition) as compared to their shuffled, surprisal-matched counterparts (SHUFFLED JABBERWOCKY, Figure 5A). This pattern follows if both systems represent aspects of syntactic structure similarly, at least in part, above and beyond next-word predictability (Figure 1, middle column). This interpretation is supported by complementary LLM probing and EEG decoding analyses which we discuss below, as well as by a body of research on language comprehension in the brain and LLMs.

The idea that syntactic structure is encoded in the brain and, to some extent, LLMs, receives independent support from each field. A rich literature on the neurobiology of syntax suggests that the brain builds hierarchical syntactic structure in time to infer meaning from sensory input [5,7,8]. Syntactic structure building operates over detailed symbolic representations [25,71,74], and goes beyond sensory, statistical, and semantic cues in the input [6,26,42,75,76]. Recently, research on LLMs suggests that syntactic structure may also be encoded in deep language models [27]. For example, LLMs are biased by hierarchical structure in a word-deletion task in a human-like manner [30]. Yet, limited studies explicitly target how the syntactic knowledge of LLMs dissociates from surface statistical patterns (but see [33]). It is also unclear the degree to which the two systems capture syntax in similar manners. Our results confirm, at least in the domain of LLM-EEG alignment, that some aspect of syntactic structure appears to be shared between these systems independent of predictability.

Our probing and decoding results are consistent with the literature reviewed above and further show that syntactic structure building in both LLMs and the brain can be decoupled from next-word predictability. LLM probing (Figure 6A) showed that most hidden layers in GPT2-XL (including layer 17 used for alignment) reliably distinguish sentences that contain syntactic structure (JABBERWOCKY) from sentences that do not (SHUFFLED JABBERWOCKY). Crucially, the sensitivity to syntactic structure in GPT2-XL is not reducible to semantic association or next-word predictability because both conditions under investigation were matched in surprisal and contained pseudowords. EEG decoding (Figure 6D) yielded parallel findings in the brain: As syntactic structure unfolds in JABBERWOCKY sentences, the scalp topography of brain activity becomes more distinguishable from that elicited by unstructured SHUFFLED JABBERWOCKY sentences. The difference in activation pattern emerges after the first few words (1-2 s after stimulus onset) and is more sustained in later portions of the sentence, after around 3 s (see, for example, [77,78]; compare also to work using parametric manipulations of phrase structure [7,42,79]). Taken together, the probing and decoding analyses show that both systems independently encode the surprisal-matched, lexically controlled contrast about syntactic structure. This convergence, in turn, accounts for the enhanced LLM-brain alignment we observe for word sequences containing syntactic structure, which is consistent with the hypothesis laid out in Figure 1 (middle column).

Our results contrast with the findings by Kauf and colleagues [11] who tested the role of syntax in alignment by perturbing syntactic structure in the materials presented to LLMs. However, this perturbation applied to LLMs, not human participants. The alignment value was instead between perturbed LLM embeddings and fMRI data from participants who only read normal unperturbed sentences. Any change in alignment in that study, or absence of difference thereof, is thus more indicative of the change in LLM internal representation *per se*, not the alignment between two systems processing the same input. The current study, on the other hand, presents syntactically manipulated materials to both comprehension systems and directly shows that GPT2-XL and human EEG activity encode syntax similarly.

The spatio-temporal profile of the syntax-induced increase in alignment is consistent with late electrophysiological signatures of syntactic composition [77,80] (Figure 5A). By averaging alignment scores over all words in a sentence-level syntactic contrast, we observed an increase in alignment spanning 650 to 1000 ms that extends over nearly all sensors, with qualitatively higher magnitudes on central-posterior sensors. In prior work, compositional effects in this time window have been reported, especially when researchers use longer sentences and narratives. For example, Li and colleagues [80] observed increased MEG activity associated with linguistic composition at around 600 ms in a naturalistic story-listening MEG dataset. When the same experimental manipulation was applied to MEG data from a controlled experiment, an earlier effect was found at around 200 ms, suggesting the influence of material design on the latency of compositional effects. Similarly on a naturalistic EEG dataset, Hale and colleagues [77] observed that parsing effort is associated with EEG activity in a 600 ms time window. The late effects in the literature and the current study may be due to the fact that syntactic operations occurred at multiple positions in our materials. The late effect we observed contrasts with studies that found earlier effects for a different, more focused set of syntactic manipulations [81–83]. Notably, these studies used contrasts that only apply to a specific point in a sentence, while our manipulation contains syntactic differences spanning across words in a sequence for two sentence templates that differed in length and structure. The increase in alignment thus reflects shared structure-building processes operating at the whole-sentence level.

The observed similarity between GPT2-XL and EEG in at least some syntactic representations must be understood within the scope of our manipulation. That is, the current study operationalized syntax as the mere presence or absence of grammatical structure for a particular set of constructions. Under this specific operationalization of syntax, LLMs do show emerging human-like ability to learn some latent structure through statistical patterns [27,30]. However, the present data do not allow us to draw conclusions about similarities for a wider range of syntactic phenomena. Indeed, the literature of human sentence processing has explored diverse aspects of syntactic computations, spanning agreement [84], long-distance dependencies [85], labeling [71], and more. Whether these separable finer components are captured in a human-like way by LLMs is still widely debated [21–23,86,87]. Taken together, our JABBERWOCKY vs. SHUFFLED JABBERWOCKY manipulation about syntax contributes to the evidence that LLMs and the brain align on a basic structure-building aspect of syntactic processing. It remains an open question whether the two systems align on more nuanced aspects of human syntax.

### 4.2. Compositional semantics is uniquely encoded in the brain

The presence of hierarchical meaning composition, or **compositional semantics**, led to a reduction in the alignment between GPT2-XL and EEG. This pattern is predicted by the hypothesis that compositional semantics is uniquely encoded in the brain (Figure 1, right column). Our findings on compositional semantics also question the “prediction-only” account [9], according to which these linguistic manipulations would not influence alignment.

In our study, the effect of compositional semantics was operationalized as the interaction between syntactic structure and lexicality. Specifically, we tested this effect through the difference-in-difference expressed as (COLORLESS GREEN - JABBERWOCKY) – (SHUFFLED COLORLESS - SHUFFLED JABBERWOCKY), which is equivalent to the joint contribution of syntax and lexicality to alignment. Compositional semantics, therefore, can be interpreted as syntax-mediated composition of real words in the current study.

When a sentence involves compositional semantics, the activation pattern of human EEG and GPT2-XL diverge, leading to a reduction in alignment (Figure 5B). Probing and decoding results indicate that such a divergence stemmed from the encoding of compositional semantics only in the brain, but not in GPT2-XL. Human EEG activity reliably distinguishes sentences containing real words vs. pseudowords (Figure 6B, 6H). Crucially, the brain shows greater sensitivity to the lexicality contrast when it involves syntactic structure, as shown in the difference in decoding accuracy between COLORLESS GREEN vs. JABBERWOCKY and SHUFFLED COLORLESS vs. SHUFFLED JABBERWOCKY. Thus, the joint contribution of syntactic structure and lexicality, or compositional semantics, is evident in EEG activity with distinctive temporal spatial patterns over the scalp. On the other hand, the interaction between syntax and lexicality does not induce additional detectable differences in LLM hidden states (Figure 6E). Although LLM hidden states reliably distinguish word sequences with real words from those with pseudowords, adding syntactic structure to those sequences does not create further representational differences that we detect. That is, there is no difference in decoding accuracy of lexicality as a function of syntax (Figure 6G). LLM probing and EEG decoding results thus converge to suggest that compositional semantics, as the joint contribution of syntactic structures and real words, is only captured by human EEG but not the hidden states of GPT2-XL. This discrepancy in processing led to a reduction in LLM-brain alignment.

The alignment reduction induced by compositional semantics was observed at 825–1000 ms over posterior sensors. This effect for compositional semantics is nested within the main effect of syntactic structure both in space and time (550–1000 ms, over all sensors; see section 3.3). This nesting is in line with compositional semantics’ dependence on syntactic structure building, which is often associated with late electrophysiological signatures as detailed above (see [78,80,88]). The interaction effect for compositional semantics effect dissociates from the main effect for lexicality observed earlier (between 50–500 ms and 650–825 ms.) This dissociation is compatible with a wide range of research on the relationship between lexical processing beginning around 200 ms and subsequent higher-level computations (see review by [89]).

The present results indicate that compositional semantics is uniquely encoded in the brain, but our understanding of compositional semantics in both systems remains understudied. Regarding its neural underpinnings, a small number of studies have attempted to characterize the neural signatures of specific semantic operations. For example, Pylkkänen and colleagues have explored the effects of specific semantic operations on MEG activity, including quantification [90] and conceptual specificity [91]. Baggio and colleagues show that different subtypes of logical-semantic operations have different ERP signatures [92,93]. Our current findings are not specific to particular semantic operations, but join other calls for more attention to this core aspect of language in the brain [94]. The scarcity of evidence also applies to research on LLMs. Although human-like understanding of individual words can be captured by static word embeddings [2,95,96], tasks that require higher semantic capacity pose a challenge even to more advanced models. For example, GPT-3, a model that is more advanced than GPT2-XL tested here, underperforms human baseline on tasks such as semantic entailment and contextual disambiguation [97]. Furthermore, direct evidence on whether they comprehend and convey complex meaning in the same rule-governed way as humans do remains elusive. Our results about compositional semantics expand on this growing literature by showing that compositional semantics is a key site of divergence between LLMs and the brain.

### 4.3. Both LLMs and the brain track associative semantics

**Associative semantics**, tested by contrasting SHUFFLED COLORLESS and SHUFFLED NORMAL sentences, does not have any significant effect on the alignment between EEG and GPT2-XL. This outcome implies the scenario in which alignment stays constant because both the LLM and the brain track semantic association similarly and uniformly across input types (Figure 1, left column).

Semantic associations play a central and well-studied role in language processing for both humans and LLMs. In humans, semantic associations underlie familiar phenomena such as semantic priming with robust behavioral effects (see [47]). Semantic associations are captured by the brain, too, such that the brain forms associations in a word sequence regardless of grammaticality [50] or deficiency in higher-level language functions such as syntactic processing [49]. In LLMs, semantic representations with human-like similarity profiles are among the first attributes to emerge as models better approximate language use [96]. Such semantic knowledge in models has been shown to systematically align with the cortical representation of word meaning [95], suggesting some similarity between the two systems regarding the representation of single-word meanings.

Our LLM probing results are consistent with prior studies and indicate that GPT2-XL is sensitive to whether words in a sentence are semantically close to each other (Figure 6C). Similarly, EEG response reliably encodes how each word semantically relates to previous words (see Supporting Information S1; also see [73]). Taken together, the absence of difference in alignment, significant LLM probing results, reliable EEG encoding of semantic associations, and well-established electrophysiological findings on semantic associations suggest that associative semantics is encoded similarly in both systems. We note that the encoding of associative semantics is not evident in EEG waveforms time-locked to the beginning of each sentence (see Figure 6F for decoding results, and Supporting Information S2 for an event-related waveform analysis). We believe this is likely due to aspects of the experimental design where the target contrast was implemented in shuffled word sequences. This design differs from typical electrophysiological studies where semantic association is at a specific word position in grammatical sentences [98] or single-word lexical designs [99]. The use of shuffled word sequences could reduce the sensitivity of a sentence-level decoder that is time-locked to sentence onset to spatial-temporal differences distributed unevenly in time. To check this, we conducted an additional TRF analysis, which is sensitive to the word-by-word evolution of linguistic features [72], and found reliable modulation of word-level EEG signals by semantic associations.

Our findings again diverge from that of Kauf et al. who highlighted the role of lexical-semantic content in LLM-brain alignment by only presenting LLMs with sentences from which the meaning content were altered or removed [11]. In that study, a drop in alignment was observed for those stimuli perturbed in meaning. Despite being different at first glance, their findings can be reconciled with our interpretation that word semantics is similarly encoded in LLMs and the brain. In Kauf and colleagues’ study, sentences with content altered or removed were presented only to LLMs, not human fMRI participants, to derive alignment measures. Since semantic content is similarly encoded in both systems, it is expected that altering lexico-semantics only in the LLMs significantly reduces alignment. Our study, on the other hand, demonstrates this possibility from a complementary perspective: Because both systems encode semantics similarly, altering the strength of association in their input causes comparable changes in both systems and thus yields no differences. In sum, the alignment between GPT2-XL and the brain stayed constant across our manipulation of associative semantics, indicating shared encoding of the association between word meanings in both systems, regardless of whether these words form syntactic structures.

### 4.4. Is next-word prediction^1^ good enough to link models and the brain?

Next-word prediction has been proposed as a parsimonious linking hypothesis between computational language models and the brain [1,9]. Results from the current study add to this view in that when next-word predictability is controlled, sentences that differ in compositional linguistic features modulate the alignment between GPT2-XL, a model believed to be highly brain-aligning, and the brain. Specifically, syntax increases alignment and compositional semantics reduces alignment. These findings strengthen the view that measures of predictability (e.g., surprisal in the current study) conflate statistical co-occurrence with other relevant features, such as hierarchical linguistic composition. Such conflation may hinder a mechanistic understanding of the computations supporting language comprehension.

Prediction is a general cognitive mechanism enabling a system to process upcoming inputs more efficiently given prior information [18,100]. In language comprehension, humans and computational models make probabilistic predictions about the next word at the phonemic, semantic, and syntactic levels [46,102–104] to handle uncertainty in the language input [34,105]. In this sense, prediction is a cognitive principle that operates in tandem with a system’s knowledge of the language being processed.

Nevertheless, we suggest caution when leveraging the notion of prediction for explaining the representations that drive LLM-brain alignment. Questionable interpretations of prediction include: (1) treating next-word prediction as the only and shared mechanism of language comprehension [9], and (2) relying solely on scalar predictability measures to link models and the brain. They share a common limitation that the mapping between the output of predictive processes and the underlying representation is underdetermined. For example, Slaats and Martin [21] demonstrated that a high predictability value (quantified as surprisal) from a language model may be consistent with either ungrammatical or unlikely utterances. Hu and colleagues [33] formalized this entanglement by showing that sentence predictability is jointly determined by both grammaticality and the likelihood of the implied meaning. Thus, the same predictability value can stem from different underlying cognitive processes (an example of the familiar problem of multiple realizability [3]).

Findings from the current study extend this critique and carry implications for using next-word prediction to explain alignment results. First, inconsistent with prediction-only accounts, surprisal-matched materials carry compositional linguistic features that are encoded by one or more systems which in turn modulate alignment. Compositional semantics is encoded more uniquely by the brain, and basic syntactic structure is encoded, at least in part, by both systems. Namely, surprisal-matched conditions induced heterogeneous rather than uniform alignment patterns. This pattern does not follow if language processing is reducible to next-word prediction. Second, our findings reveal a limitation of relying solely on scalar measures of predictability. Both systems encode basic syntactic structures, yet model-derived surprisal does not distinguish sequences with and without those structures, nor does it capture either system’s sensitivity to that distinction or the alignment between systems. Therefore, even though next-word predictability is useful for modeling behavioral and neural data [106–108], it cannot alone replace mechanistic explanations in explaining LLM-brain alignment.

Two possible paths emerge to better advance the development of such mechanistic accounts. The first approach is to consider processing components that act alongside predictive processing. Examples of this approach are studies testing how linguistic reanalysis plays a crucial role in explaining human language comprehension above and beyond next-word prediction [22,87,109]. On this view, reanalysis is a cognitive process that operates externally to the predictive architecture, capturing variance that predictability alone cannot.

Another direction forward is to emphasize mechanistic explanations in predictive systems rather than relying on the predictability readout [17,76,110]. Take syntax as an example: Basic syntactic computations have been proposed and tested both in the neurobiology of language [8,25,68] and computational language modeling [27,30,111]. Crucially, identifying individual mechanistic components specifies, rather than undermines, the predictive architecture of language processing both in the brain [17,25,76] and models [27,30,76]. Thus, insights into the underlying computations provide additional, not competing, evidence for how a language processing system generates predictions based on interpretable latent knowledge.

## 5. Conclusions

Emerging findings of LLM-brain alignment on large naturalistic datasets have been taken to support the claim that the two systems may operate in a similar way. In this study, we test this inference with a more systematic approach. By manipulating syntax, compositional semantics, and associative semantics in the input to both systems, we reveal the components of language on which LLMs and the brain align and misalign: Basic syntactic structure and semantic association are encoded similarly in both systems, whereas compositional semantics is more uniquely encoded in the brain. These findings are incompatible with the “prediction-only” account of LLM-brain alignment in as much as hierarchical linguistic composition modulates alignment when predictability is tightly controlled. To conclude, the current study emphasizes the importance of hierarchical compositional linguistic structure in understanding both biological and artificial language processing systems.

## Acknowledgments

We thank Jingzhu Peng and Quan Zou for their help with data collection. We thank members of the Computational Linguistics Lab and the Psycholinguistics Discussion Group at the University of Michigan for discussions about this project.

## Footnotes

1 In this section we distinguish three related concepts: prediction, next-word prediction, and predictability. Prediction refers to the general cognitive mechanism that applies to language [18] and other domains of cognition [100]. Next-word prediction refers specifically to the step in language comprehension at which the brain or a computational language model generates a probability distribution for the upcoming word. A processing system may, but does not necessarily, possess mechanistic knowledge to generate next-word predictions [9,101]. Predictability refers to the scalar measure derived from the process of next-word prediction [34].

