## Supporting information for "Beyond next-word prediction: hierarchical linguistic composition modulates LLM-brain alignment in time"

**S1 text. Temporal response function analysis of semantic dissimilarity**

Our manipulation of semantic association was implemented across shuffled word sequences instead of at a specific word position in grammatical sentences; the sensitivity of a time-locked sentence-level decoder may be compromised in this circumstance. To better accommodate temporally scattered responses to associative semantics, we conducted a temporal response function (TRF) analysis to test whether EEG tracks the word-by-word pattern of associative semantics.

The TRF analysis on EEG involves estimating a temporal kernel, which, when convoluted with a linguistic feature (as a predictor), explains the unique EEG response to that feature [66]. For our purpose, we estimated TRFs to semantic dissimilarity which quantifies the semantic distance between the current word and its preceding context. Semantic dissimilarity was calculated as

$$D(w_i) = 1 - \frac{v_{w_i} \cdot \bar{v}_{1:i-1}}{||v_{w_i}||\ ||\bar{v}_{1:i-1}||}$$

where the fraction is the cosine similarity between the GloVe embedding of the current word and the average embedding of all preceding words in a sentence. This predictor is constructed following earlier work by Broderick and colleagues [73].

To test whether semantic dissimilarity explains additional variance in EEG, we estimated and compared two TRF models: The base model includes the following predictors at each word: word length, word onset (binary), word frequency (based on SUBTLEX-US [55]), and GPT2-XL surprisal. The based model is compared with another model that additionally includes semantic dissimilarity as a predictor. TRF models were estimated for each condition (SHUFFLED NORMAL, SHUFFLED COLORLESS) and each participant using the `boosting` function in the *eelbrain* python package (version 0.39.0) [66], with *l*2 error, 50-ms hamming windows, 4-fold cross-validation, and early stopping. The proportion of variance explained were then extracted from the base and full TRF models and compared using a spatial cluster-based permutation test.

Semantic dissimilarity explains more variance than base predictors in both SHUFFLED NORMAL and SHUFFLED COLORLESS sentences, indicating that the word-by-word evolution of semantic association is meaningfully encoded in the brain. SHUFFLED NORMAL: $t(27) = 9.81$, $p < 0.001$. SHUFFLED COLORLESS: $t(27) = 9.90$, $p < 0.001$. There was no significant difference in additional variance explained by semantic dissimilarity between the two conditions. $t(27) = -0.01$, $p = 0.94$.

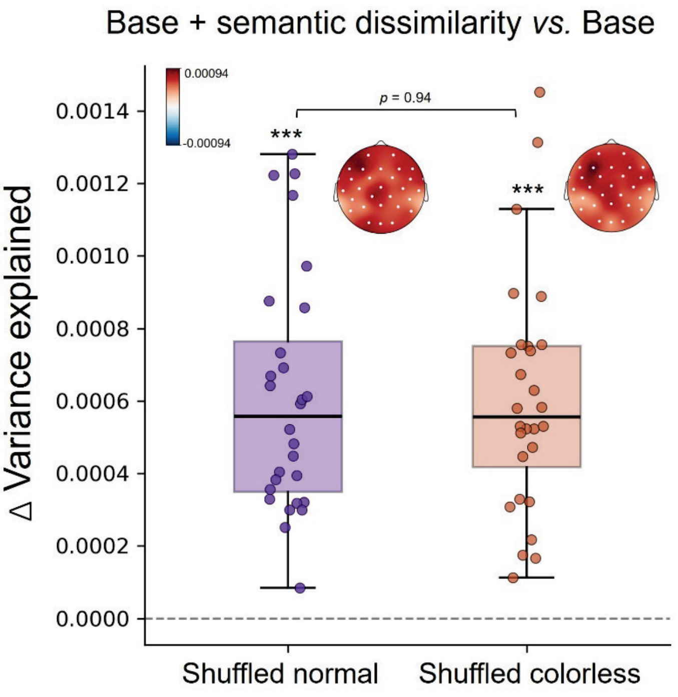


**Figure S2. Semantic association explains additional variance in EEG response.** Additional variance explained is computed by subtracting the variance explained by base predictors from the variance explained by base and semantic dissimilarity predictors. Topomap insets illustrate the scalp distribution of the additional variance explained. White dots represent sensors that show a significant increase in variance explained.

**S2 text. Time-locked waveform analysis of associative semantics**

We compared the EEG waveforms in SHUFFLED NORMAL (high semantic association) and SHUFFLED COLORLESS (low semantic association) conditions. Both 5- and 8-word long sentences were included in the analysis. EEG data were filtered to 0.1-20 Hz, epoched from -0.3 to 3 seconds, baseline-corrected to -0.3 to 0 seconds, and averaged across participants. This time window covers the initial five words. A spatial-temporal permutation cluster *t*-test did not find a significant difference in any spatial-temporal cluster.

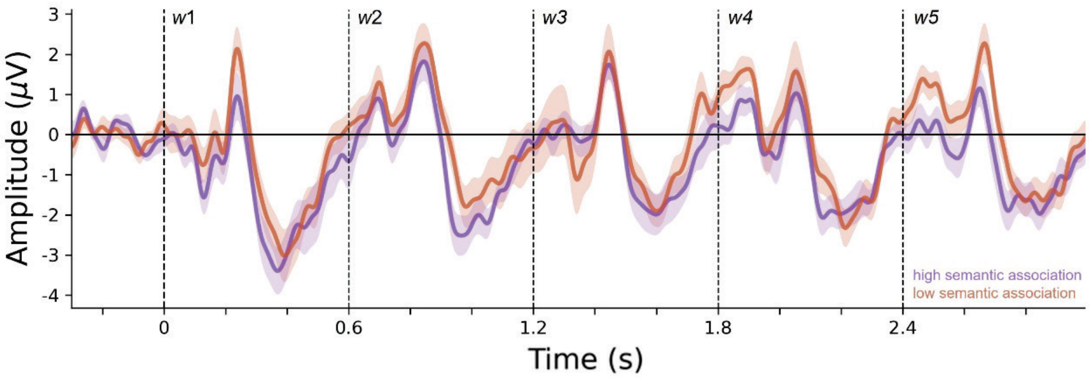


**Figure S2. EEG waveform comparison for the contrast for associative semantics (SHUFFLED NORMAL and SHUFFLED COLORLESS).** EEG data were averaged across central-posterior sensors (Cz, C3, C4, CP1, CP2, Pz, P3, P4) for illustration. Shaded areas indicate one standard error of the mean.
